# A naturalistic environment to study natural social behaviors and cognitive tasks in freely moving monkeys

**DOI:** 10.1101/2020.09.25.311555

**Authors:** Georgin Jacob, Harish Katti, Thomas Cherian, Jhilik Das, Zhivago KA, SP Arun

**Author notes:** indicate authors with equal contributions.

## Abstract

Macaque monkeys are widely used to study the neural basis of cognition. In the traditional approach, the monkey is brought into a lab to perform tasks while it is restrained to obtain stable gaze tracking and neural recordings. This unnatural setting prevents studying brain activity during natural, social and complex behaviors. Here, we designed a naturalistic environment with an integrated behavioral workstation that enables complex task training with viable gaze tracking in freely moving monkeys. We used this facility to train monkeys on a challenging same-different task. Remarkably, this facility enabled a naïve monkey to learn the task merely by observing trained monkeys. This social training was faster primarily because the naïve monkey first learned the task structure and then the same-different rule. We propose that such hybrid environments can be used to study brain activity during natural behaviors as well as during controlled cognitive tasks.

## INTRODUCTION

> “Thus it is said:
>
> The path into the light seems dark,
>
> the path forward seems to go back”
>
> — -- Tao Te Ching v41 (Mitchell, 1988)

Macaque monkeys are highly intelligent social animals, because of which they are widely used to understand the neural basis of cognition (Passingham, 2009). Insights from primate studies have yielded important insights into brain disorders (Roelfsema and Treue, 2014; Buffalo et al., 2019). In the widely used traditional approach, brain activity is recorded from monkeys while their head is restrained to obtain stable eye tracking and wired neural recordings. This highly unnatural setting blinds us from studying brain activity “in the wild” while animals engage in natural, social and complex behaviors. Despite these limitations, the traditional approach is still widely used since it remains the only viable way to obtain stable gaze and record neural signals in monkeys performing cognitive tasks.

Recording brain activity during natural and complex behaviors requires overcoming several major challenges. First, it should be possible to record neural activity wirelessly so that animals can move freely. This has recently become possible with the advent of wireless technologies (Chestek et al., 2009; Szuts et al., 2011; Roy and Wang, 2012; Berger et al., 2020). Second, animals must be housed in a hybrid environment for them to repeatedly engage in complex cognitive tasks for rigorous studies of behavior, as well as engage in interesting natural and social behaviors in their home environs, with the flexibility to go back and forth freely between them. The design principles for such naturalistic environments as well as standard procedures to maximize animal welfare are well understood now (Woolverton et al., 1989; Röder and Timmermans, 2002; Honess and Marin, 2006; Seier et al., 2011; Cannon et al., 2016; Coleman and Novak, 2017). Recent studies have demonstrated that monkeys can be trained to perform complex tasks using touchscreen devices that can be easily integrated into a naturalistic environment (Rumbaugh et al., 1989; Mandell and Sackett, 2008; Fagot and Paleressompoulle, 2009; Gazes et al., 2013; Calapai et al., 2017; Claidière et al., 2017; Tulip et al., 2017; Berger et al., 2018). Finally, it should be possible to obtain stable gaze signals from unrestrained animals. This is a non-trivial challenge because most infrared eye trackers require restraining the head (Machado and Nelson, 2011), and do not work at close quarters with an elevated line of sight as would be the case for a monkey interacting with a touchscreen.

Here, we designed a hybrid naturalistic environment with a touchscreen workstation that can be used to record brain activity in controlled cognitive tasks as well as during natural and social behaviors. We show that this facility can be used to obtain viable gaze signals and that it can be used to train monkeys on a same-different task by taking them through a sequence of subtasks with increasing complexity. Remarkably, we show that a naïve monkey was able to learn the entire task much faster merely by observing two other trained monkeys.

## RESULTS

### Facility overview

We designed a novel naturalistic environment for recording brain activity during controlled cognitive tasks as well as natural and social behaviors (Figure 1). Monkeys were group-housed in an enriched living environment with access to a touchscreen workstation where they could perform cognitive tasks for juice reward while their brain activity is recorded (Figure 1A; see Methods). The enriched environment comprised log perches and dead trees with natural as well as artificial lighting with several CCTV cameras to monitor movements (Figure 1B). We also included tall perches to which animals could retreat to safety (Figure 1C). The continuous camera recordings enabled us to reconstruct activity maps of the animals with and without human interactions (Figure 1D; [Video S1](https://osf.io/3kn7m/)). To allow specific animals access to the behavior room, we designed a corridor with movable partitions so that the selected animal could be induced to enter while restricting others (Figure 1E). We included a squeeze partition that was not used for training but was used if required for administering drugs or for routine blood testing (Figure 1F). This squeeze partition had a ratchet mechanism and locks for easy operation (Figure 1G). After traversing the corridor (Figure 1H), monkeys entered a behavior room containing a touchscreen workstation (Figure 1I). The behavior room contained copper-sandwiched high pressure laminated panels that formed a closed circuit for removing external noise (Section S1). The entire workflow was designed so that experimenters would never have to directly handle or contact the animals during training. Even though the facility contained safe perches out of reach from humans, we nonetheless developed standard protocols to easily isolate each monkey and give it access to the behaviour room (see Methods).

**Figure 1:**
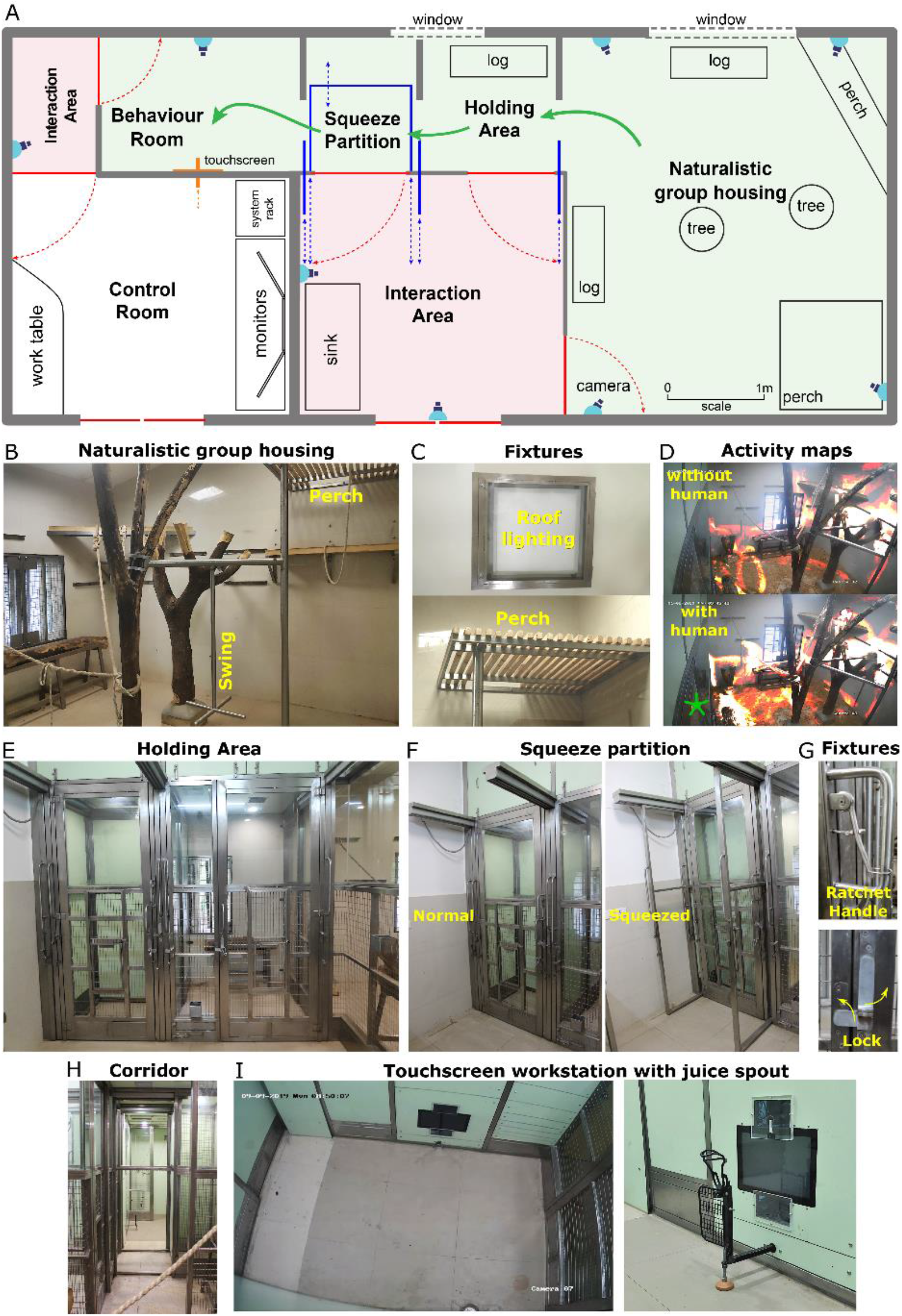
Hybrid naturalistic environment with touchscreen workstation. (A) Illustrated layout of the facility designed to enable easy access for monkeys to behavioral tasks. Major features placed for enrichment are labelled. *Blue lines* indicate partitions for providing access to various portions of the play area. Typical movement of an animal is indicated using *green arrows. Red lines* indicate doors that are normally kept closed. (B) View into the play area from the interaction room showing the enriched environment. (C) *Top*: Close up of the perch that provides monkeys with an elevated point of observation. *Bottom*: Roof lights that have been enclosed in stainless steel and toughened glass case to be tamper-proof. (D) *Top*: Heatmap of residence duration of monkeys (red to yellow to white = less to more time spent in location) in the play area analyzed from a ~7 min video feed of the CCTV in panel I There was no human presence in the interaction room during this period. *Bottom*: The same residence analysis but was human presence in the interaction room during a ~7 min period on the same day. (E) View from below the CCTV in the interaction area to the squeeze and holding areas with trap-doors available to bring monkey into chairs. (F) The squeeze room constricted for restraining monkeys within. *Left*: View of the room in a normal condition *Right*: View of the room in the squeezed condition. (G) *Top*: Close view of the rachet to bring the squeeze partition forward. *Bottom*: Close view of the partition lock. (H) View of the path taken by monkey from play area through the holding and squeeze area into the behavior room. **(I)** Top-down view from the CCTV in the behavior room showing the placement of the touchscreen on the modular panel wall and the abutting juice reward arm in from of it.

### Touchscreen workstation with eye tracking in unrestrained monkeys

The touchscreen workstation is detailed in Figure 2. Monkeys were trained to sit comfortably at the juice spout and perform tasks on the touchscreen for juice reward. The workstation contained several critical design elements that enabled behavioral control and eye tracking, as summarized below.

**Figure 2:**
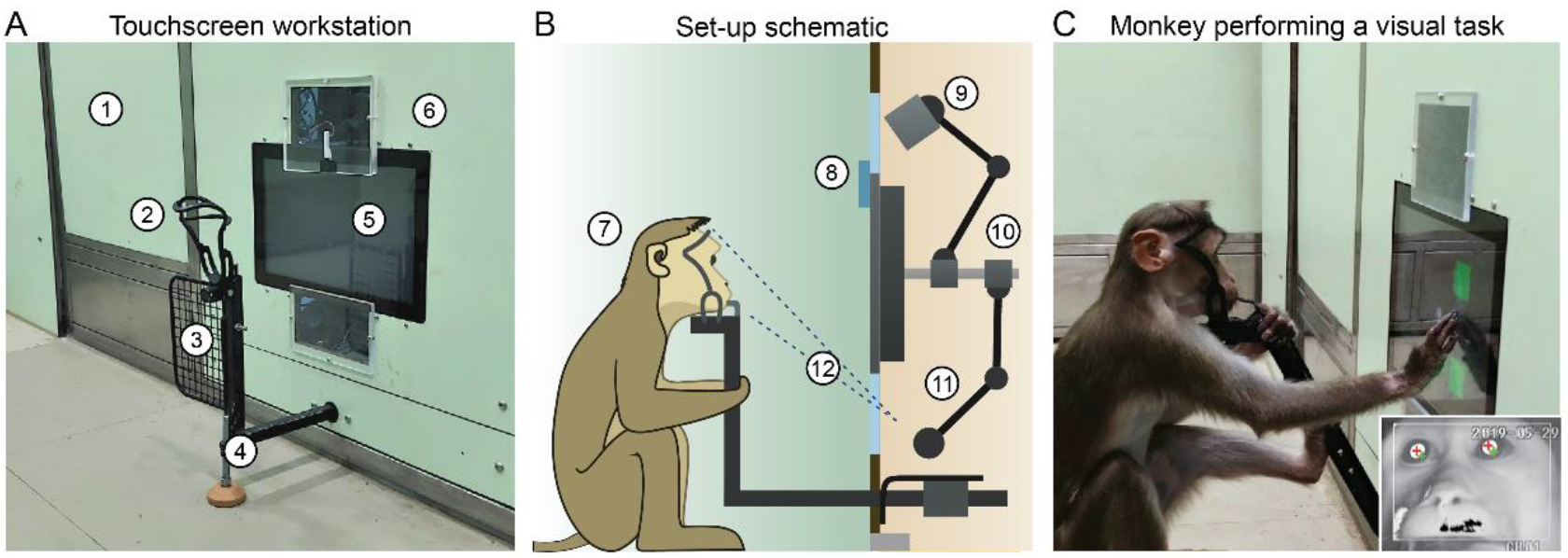
Touchscreen workstation with eye tracking for unrestrained monkeys. *(A)* Labelled photograph of the touchscreen workstation from the monkey’s side. *Labels: 1: Partition panel with electromagnetic shielding; 2: Whole head frame; 3: Grill to block left hand access; 4: Movable reward delivery arm with concealed juice pipe; 5: Touchscreen; 6: Transparent viewports*. *(B)* Labelled cross section showing both monkey and experimenter sides. *Labels: 7: Position of monkey at workstation; 8: Channel for mounting photodiode; 9: Synchronized optical video camera; 10: Adjustable arms mounted on shaft behind touchscreen back panel; 11: Eye tracker infrared (IR) camera with IR illuminator; 12: Field of view of the eye tracker*. (C) Photograph of monkey performing a task. Note that the reward arm shown is an older version of reward arm shown in (A). *Inset*: Thresholded image from the infrared eye tracker of the monkey, showing the detected pupil (white disk) with cross-hairs representing the detected pupil center and corneal reflection.

First, we developed a juice delivery arm with a drain mechanism that would take any extra juice back out to a juice reservoir (Section S1). This was done to ensure that monkeys drank juice directly from the juice spout after a correct trial instead of subverting it and the accessing the spillover. Second, we developed several modular head frames that were tailored to the typical monkey head shape (Figure 2B; see also Section S1). In practice, monkeys comfortably rested their chin/head against these frames and were willing to perform hundreds of trials even while using the most restrictive frames. Third, we included a removable hand grill to prevent the monkeys from accessing the touchscreen with the left hand (Figure 2A). This was critical not only for reducing movement variability, but also to enable a direct line of sight for the eye tracker. Finally, we affixed two transparent viewports above and below the touchscreen, one for an optical video camera and the other for an infrared eye tracker (Figure 2A-B). This design essentially stereotyped the position of the monkey’s head and gave us excellent pupil and eye images (Figure 2C, inset) and consequently good eye tracking.

### Same-different task with gaze-contingent eye tracking

Understanding the neural basis of cognition often requires training monkeys on complex cognitive tasks with events contingent on their eye movements, such as requiring them to fixate. As a proof of concept, we trained two animals (M1 & M3) on a same-different (or delayed match-to-sample) task with real-time gaze-contingency.

The timeline of the task is depicted schematically in Figure 3A. Each trial began with a hold cue that was displayed until the animal touched it with his hand, after which a fixation cross appeared at the center of the screen. The monkey had to keep its hand on the hold cue and maintain its gaze within a 3° window around the fixation cross from then on. Following this a sample image appeared for 500 ms after which the screen went blank for 200 ms. After this, several events happened simultaneously: a test stimulus appeared, the hold cue disappeared, fixation/hold conditions were removed and two choice buttons appeared above and below the hold cue. The animal had to make a response by touching one of the choice buttons within 5 s. The test stimulus was presented till the monkey makes a response. If the test image was identical to the sample, the monkey had to touch the upper button or if it was different, the lower button. Example videos of the same-different task and a more complex part-matching task are shown in [Video S2](https://osf.io/ftgjy/).

**Figure 3:**
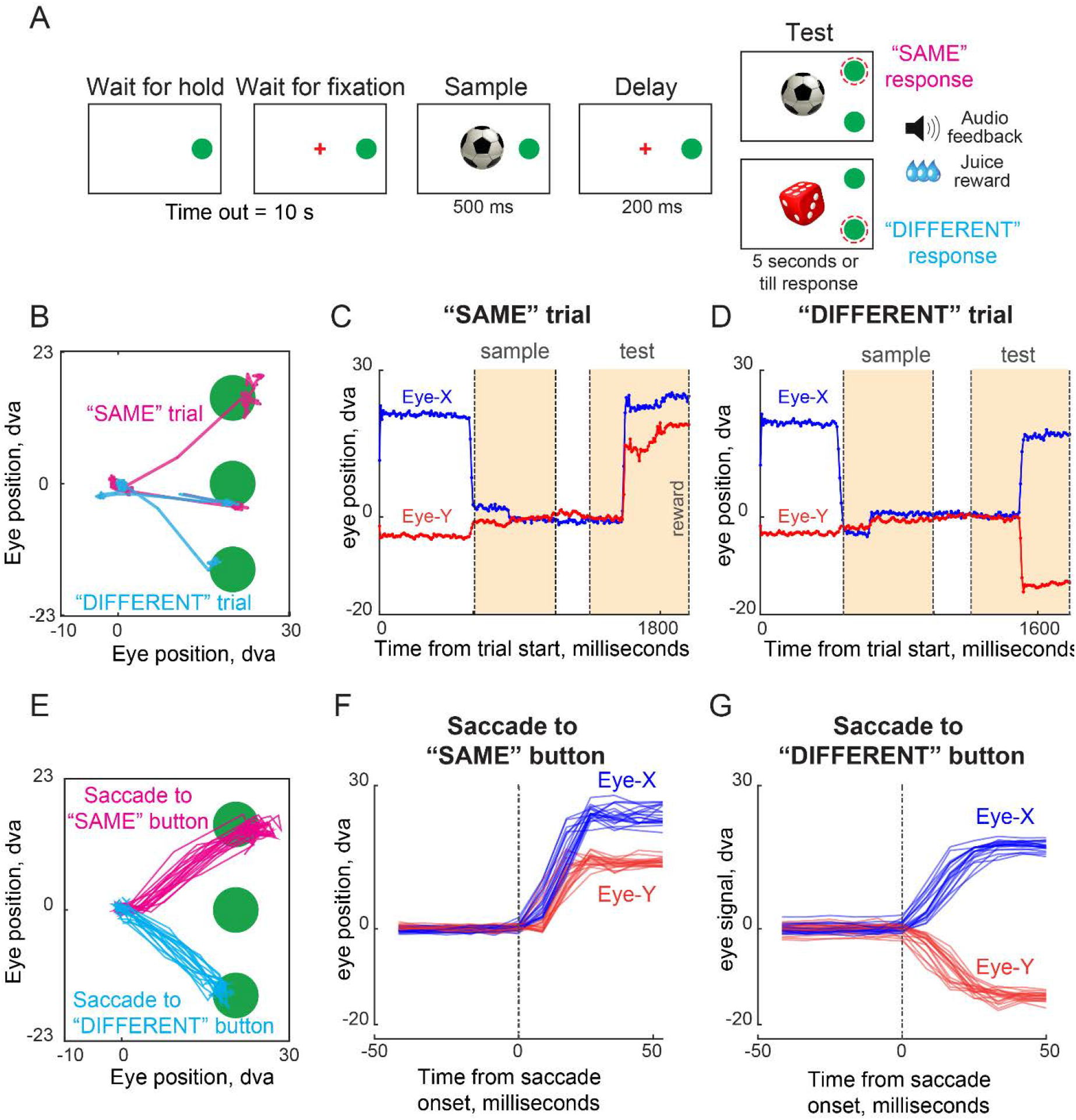
Same Different Task with gaze-contingent eye-tracking. (A) The diagram shows the sequence of stimulus events and expected monkey behaviour during a single trial of same task. (B) Eye traces overlaid on the stimulus screen, for one representative same trial (magenta) and one representative different trial (blue). The monkey initiated a trial by touching and holding the green middle response button following which a central fixation cue was presented. When the monkey looks at the central fixation cue, first a sample stimulus is shown followed by a test stimulus after a brief delay. The monkey had to touch the upper/lower green button if the test did/did not match the sample. (C) Time series eye gaze data obtained for horizontal (blue) and vertical (red) components of eye movements made by the monkey for the SAME trial shown in (B). Dotted lines mark onset of the sample image, test image and reward delivery. (D) Time series eye gaze data shown for a correct DIFFERENT choice trial in (B). (E) Gaze samples during saccades made towards response buttons during SAME response trials *(magenta)* and DIFFERENT response trials *(blue)*. (F) Eye position traces as a function of time (aligned to saccade onset) for the SAME response trials shown in (E). Saccade onset was defined based on the time at which saccade velocity attained 10% of the maximum eye velocity. **(G)** Same as (F) but for DIFFERENT response trials.

Figure 3B illustrates the example gaze signals recorded from monkey M1 during two trials of the same-different task, one with a “SAME” response and the other with a “DIFFERENT” response. The monkey initially looked at the hold, then at the sample, and eventually at the choice buttons. The time course of the two trials reveals eye movements in the expected directions: for the “SAME” trial, the vertical eye position moves up shortly after the test stimulus appeared (Figure 3C) whereas in a “DIFFERENT” response trial, the vertical position moves down (Figure 3D). Remarkably, despite being unrestrained, we obtained highly reliable gaze position across trials (Figure 3E), allowing us to reconstruct the characteristic time course of saccades (Figure 3F-G). This high fidelity of gaze data was largely due to the stereotyped position of the juice spout, which made the animal put its head in exactly the same position each time, enabling accurate eye tracking (see [Video S3](https://osf.io/pj57n/) for eye movements in a single trial, [Video S4](https://osf.io/pgh95/) for eye movements across trials).

### Tailored Automated Training (TAT) on same-different task

Here we describe our novel approach to training animals on this same-different task, which we term as “Tailored Automated Training” (TAT). In the traditional paradigm, before any task training can be started, monkeys have to be gradually acclimatized to entering specialized monkey chairs that block them from access to their head, and to having their head immobilized using headposts for the purpose of gaze tracking. This process can take a few months and therefore is a major bottleneck in training (Fernström et al., 2009; Slater et al., 2016; Mason et al., 2019). We bypassed these time-consuming steps in our setup, which allowed us to focus entirely on task-relevant training.

We trained two monkeys (M1 & M3) using TAT (for details, see Section S2). The well-known principle behind training monkeys on complex tasks is to take the animal through several stages of gradual training so that at every stage the animal is performing above chance, while at the same time learning continuously. On each session, we gave access to each monkey individually by separating it from its group using the holding areas (Figure 1A). Each monkey was guided automatically through increasingly complex stages of interaction with the touchscreen setup. These stages went from a basic task where the monkey received a reward for touching/holding a target square on the screen, to the full same-different task described in the previous sections. Importantly, each monkey went through a unique trajectory of learning that was tailored to its competence on each stage. There were a total of 10 stages and multiple levels within each stage. Only one task-related parameter was varied across levels in a given stage. Monkeys would progress to the next level once it completed 50 trials with at least 80% accuracy. By the end of training, both monkeys were highly accurate on the same-different task (91% for M1, 82% for M3). The duration of training from completely naïve to fully trained was approximately 90 sessions or days. Thus, the tailored automated training (TAT) paradigm deployed in this hybrid environment can enable automated training of monkeys on complex cognitive tasks while at the same time maximizing animal welfare.

### Can a naïve monkey learn the task by observing trained monkeys?

Our novel facility has the provision to allow multiple monkeys to freely move and access the touchscreen workstation. We therefore wondered whether a naïve monkey could learn the same-different task by observing trained monkeys. This would further obviate the need for the TAT paradigm by allowing monkeys to learn from each other.

To investigate this possibility, we ran social training sessions in which a naïve monkey (M2) was introduced along with a trained monkey into the behavior room. Importantly, the naïve monkey (M2) was intermediate in its social rank, with one of the trained monkeys (M1) being above it and the other (M3) being below it. On each day or training session, M2 participated in two social training sessions: one where it was introduced with M1 and the other when it was introduced with M3.

The outcomes are summarized in Figure 4. To summarize, M2 began to perform the same-different task at chance after 13 social training sessions, by which point there were no interactions between M1 & M2 (with M1 dominating throughout) and no interactions between M2 & M3 (with M2 dominating throughout). We therefore stopped social sessions and started introducing M2 by himself into the behaviour room. From here on, M2 took 8 more sessions to perform above-chance, and by the end of 29 sessions, had achieved 91% accuracy on the task. Remarkably, M2 learned the task much faster using social observation and learning than M1 & M3 did using the TAT paradigm.

**Figure 4:**
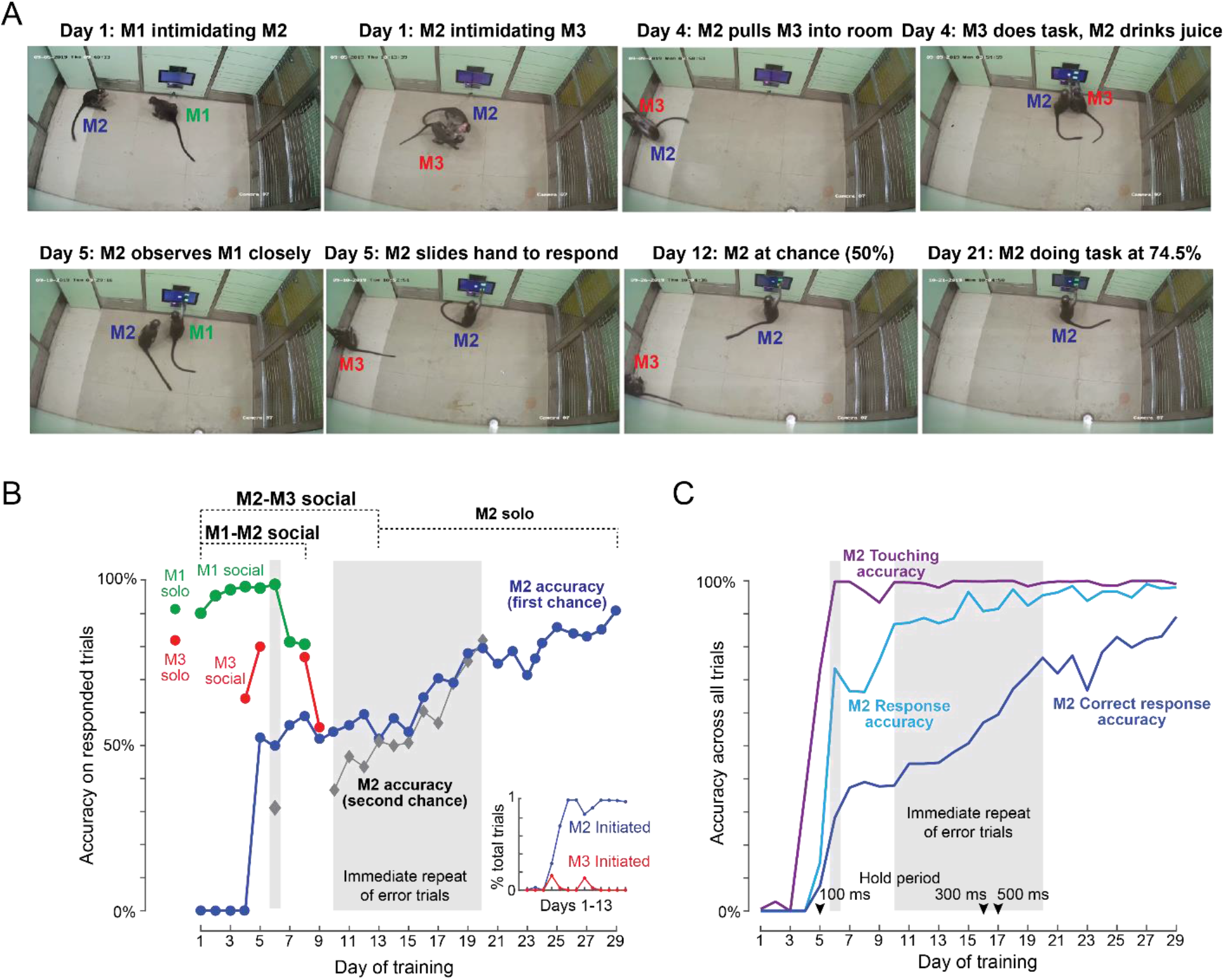
Naïve monkey learning by observing trained monkeys. A naïve monkey (M2) learned to perform the same-different task by observing two trained monkeys (M1 & M3). The social rank order is M1 > M2 > M3. See [Video S5](https://osf.io/udpbg/). (A) Surveillance camera photos depicting the important stages of learning for M2. (B) Accuracy in social training sessions (green-M1, blue-M2 and red-M3) across days. For each monkey, accuracy is calculated on trials on which it made a response (Errors due to touch failure, hold failure and non-response are excluded). Shaded regions depict days on which error trials were repeated immediately without a delay, allowing monkeys to switch their response upon making an error. M2 accuracy on such repeated trials is shown separately (*grey*). M1 and M3 accuracy on the same image set prior to and during social sessions is shown by *red* and *green* dots at the left (M1 solo accuracy: 91%, M3 solo accuracy: 82%). *Inset*: Percentage of all trials initiated by M2 (*blue*) and M3 (*red*) during M2-M3 sessions across thirteen days of training. **(C)** Accuracy for monkey M2 for various types of response, calculated as percentage of all trials. *Touching accuracy (purple)*: percentage of all trials initiated by touching the hold button. *Response accuracy (cyan)*: percentage of trials where M2 touched any choice button out of all trials. *Correct accuracy (blue)*: Percentage of trials where M2 touched the correct choice button out of all trials. Shaded regions depict days on which error trials were repeated immediately without a delay.

### Stages of social learning

How did M2 learn the task? Were there any key stages during its learning? To answer these questions, we closely inspected the CCTV videos as well as the automated training performance of all monkeys during this period. These stages are summarized in Figure 4 and selected videos of these stages are shown in [Video S5](https://osf.io/udpbg/). Trials attempted by M1, M2 and M3 are depicted in Section S3.

On Day 1, we observed interactions expected from their social rank. M1 intimidated M2 whenever M2 approached the workstation and M2 in turn intimidated M3 and prevented M3 from approaching the workstation (Figure 4A). From Day 2, M2 began touching the screen (Figure 4C, touching accuracy), but still did not get juice reward.

On Day 4, interestingly, M2 pulled M3 from the holding room into the behaviour room, apparently in an attempt to make M3 do the task. Like in previous sessions, M2 positioned himself in front of the reward tube, but interestingly, M3 approached the touch screen from one side and performed a few trials while M2 received the juice (Figure 4A). After this interaction, M2 initiated more trials by touching the hold button (Section S3) but still did not earn any juice reward by himself.

On Day 5, in the M1-M2 session, M2 watched M1 closely for long stretches. In the M2-M3 session on the same day, for a few trials, M2 maintained hold till the choice buttons appeared and ended up touching the lower button (corresponding to a DIFFERENT response) by dragging his hand down. M2 made 4 correct responses in this manner and received juice reward. For a short stretch of trials M2 allowed M3 to do the task (same as in Day 4) and both M2 and M3 shared the reward tube. For this stretch M2 received reward at a much higher rate (8 out of 14 trials, Section S3). After this M2 did not allow M3 to do any trials for rest of the session, but his response accuracy and correct response accuracy (by dragging hand through the DIFFERENT response button) increased. On this day, first chance accuracy of M2 was 53% on responded trials (Figure 4B), though this was still a small proportion of all trials (7.6%, Figure 4C).

On Day 6, in the M2-M3 session, M2 started responding on more than 70% of the trials and started making the SAME response as well once we began immediate repeat of error trials (see Methods). By Day 8, we ended M1-M2 sessions because we observed little or no interactions between them.

On Days 8 & 9, M2 allowed M3 to perform a few trials (as before) and they shared the reward, though we did not see any change in its performance after those interactions.

From Day 9 onwards, M2 did not allow M3 to attempt any more trials, while his performance hovered around chance (Figure 4B). After 4 such days, we stopped M2-M3 sessions and began to introduce M2 by himself into the behaviour room (Day 14 onwards; Figure 4B). The M2-M3 interactions are summarized in Figure 4B *(inset)*.

From Days 14-29, M2 was trained alone and learned the task by trial and error. We included an immediate repeat of error trials (Day 6 & Day 10-20), which allowed M2 to switch his response to the other choice button upon making an error. However his accuracy on both the first-chance trials (i.e. trials without an error on the preceding trial) and on second-chance trials (i.e. on trials with an error on the preceding trial) both increased monotonically, suggesting that he was continuously learning the concept of same-different and not just learning to switch on making an error (Figure 4B). By Day 25, M2 had attained an accuracy of 86%, meaning that he had learned the image same-different task.

The above observations demonstrate that M2 learned the task in two distinct phases. In the first phase, he learned the basic structure of the task through social interactions and learning – this included maintaining hold until the test image appeared and touching the one of the choice buttons subsequently. By the end of this stage he was no longer socially interacting or learning from the other two monkeys. In the second phase, he learned the same-different rule all by himself through trial-and-error. Thus, the social sessions naturally dissociated these two stages of learning, enabling faster learning compared to the more artificial stages introduced by us.

## DISCUSSION

Here, we designed a novel hybrid naturalistic environment with a touchscreen workstation that can be used to record brain activity in controlled cognitive tasks as well as during natural and social behaviors. We demonstrate two major outcomes using this facility. First, we show that viable gaze tracking can be achieved in unrestrained, freely moving monkeys working at the touchscreen on a complex cognitive task (same-different task). Second, we show that a naïve monkey could learn the same-different task much faster by socially observing two trained monkeys doing the task. We discuss these advances in relation to the existing literature below.

### Relation to other primate training facilities

Our novel hybrid environment with a touchscreen is similar to other efforts (Calapai et al., 2017; Tulip et al., 2017; Berger et al., 2018), where the common goal is a seamless behavior station to enable training monkeys within their living environment. However it is unique and novel in several respects. First, we were able to achieve precise monitoring of gaze in unrestrained animals. This is an important advance since such gaze signals are required for most complex cognitive tasks. We accomplished this by designing a juice spout with a chin and head frames that essentially enabled monkeys to achieve a highly stereotyped head position while performing the task. Second, unlike other facilities where the touchscreen workstation is an add-on or housed in a separate enclosure (Evans et al., 2008; Mandell and Sackett, 2008; Fagot and Paleressompoulle, 2009; Fagot and Bonté, 2010; Calapai et al., 2017; Claidière et al., 2017; Walker et al., 2019), our touchscreen is mounted flush onto a modular wall (with provision for expansion) that provided viewing access to other monkeys to observe the task, which enables novel social interactions such as the ones we have observed. Third, we demonstrate that monkeys can be group-housed even with safe perches out of reach from humans, yet it is possible to isolate each animal individually and give it access to the touchscreen workstation (see Methods). Finally, our behaviour room is electromagnetically shielded and therefore capable of wireless neural recordings in contrast to previously reported facilities that were largely developed only for task training and not recording.

### Social learning of complex tasks in monkeys

We have found that a naïve monkey can learn the complex same-different task much faster by observing other trained monkeys. This is a novel and exciting outcome that could signal a paradigm shift in monkey training. An extreme interpretation of this finding is that only one animal needs to be trained through TAT and other animals can learn from it through social observation, thereby bringing huge savings in human efforts. However, we caution that this approach might not work on every individual, since the learning could depend on individual ability and on relative social rank. Our finding that naïve animals can learn from experts is consistent with reports of observational learning in monkeys (Brosnan and de Waal, 2004; Subiaul et al., 2004; Meunier et al., 2007; Falcone et al., 2012; Monfardini et al., 2012). However, in these studies, naïve animals learned relatively simple tasks and did not have unconstrained access to the expert animal to observe or intervene at will. By contrast, we show that a naïve monkey can learn a cognitively demanding same-different task by unconstrained observation of trained animals.

Our social learning results offer interesting insights into how animals learn complex cognitive tasks. Learning occurred naturally in two distinct stages. In the first stage, the naïve monkey learned the basic task structure by socially observing trained monkeys, but did not learn the same-different rule. This stage took only a few days during social learning whereas we estimate it could take several weeks for us to achieve a similar outcome through a process such as TAT. In the second stage, the naïve monkey went solo, showing no interest in social observation, and learned the same-different rule by trial-and-error. This stage took about two weeks, and we estimate it would take us a similar amount of time using a process such as TAT. Thus, the major advantage of social learning was that it enabled the naïve animal to learn the basic task structure from a conspecific, while learning the more complex cognitive rule by itself.

In sum, our facility represents an important first step in turning the traditional paradigm on its head: rather than bringing the monkey into a specialized lab for brain recordings and gaze tracking, we show that it is possible to bring a specialized lab into the monkey’s home for the study of complex cognitive tasks as well as natural and social behaviors.

## METHODS

All procedures were in accordance to an experimental protocol approved by the Institutional Animal Ethics Committee of the Indian Institute of Science and by the Committee for the Purpose of Control and Supervision of Experiments on Animals, Government of India.

### Animals

Three bonnet macaque monkeys (*macaca radiata*, laboratory designations: Di, Ju, Co; all male; aged ~6 years – denoted as M1, M2, M3 respectively) were used in the study.

### Design and implementation of hybrid naturalistic facility

Our goal was to design and construct a novel facility with an enriched living environment, controlled access to a behavior room with a touchscreen workstation, and provision for training on complex cognitive tasks and eventual wireless recording of brain signals.

In most primate facilities where macaques have freedom of movement while interacting with behavior stations, the key differences typically lie in the placement of the behavior station about the living room, mode of interaction while monkeys performed tasks and the degree to which the animal’s behavior could be observed. The simpler and more common approach has been to install the behavior station directly in the living room either on the walls (Rumbaugh et al., 1989; Crofts et al., 1999; Truppa et al., 2010; Gazes et al., 2013; Tulip et al., 2017; Butler and Kennerley, 2019) or in an adjacent enclosure where a single subject can be temporarily isolated (Evans et al., 2008; Mandell and Sackett, 2008; Fagot and Paleressompoulle, 2009; Fagot and Bonté, 2010; Calapai et al., 2017; Claidière et al., 2017; Walker et al., 2019). Although the former approach is easiest to implement and can let multiple monkeys interact with the behavior station, it can be challenging to prevent a monkey from getting distracted from other events in its living environment and to ensure access for a specific monkey. In contrast, the latter approach is better suited to control for disturbances in the living room but with the caveat that it has commonly been designed for use by one monkey at a time and thus precludes studying interesting behaviors where multiple monkeys can observe or interact with the behavior station.

Here, we combined the best of both approaches such that monkeys traversed away from the living room to a behavior room that had a single behavior station mounted flush to its walls. As a result, monkeys did not require human intervention to get access to the behavior station and were thus free to come to the behavior station when they wished and interact with it for juice rewards. Our approach also enabled the dynamics between each monkey to play out and determine the duration of access by each monkey.

Such an arrangement was of benefit to us as the experimenters as well. First, the need to transfer monkeys manually to the behavior room and back using monkey chairs was avoided. Second, as the monkey would interact with the behavior station in a behavior room that was separated by multiple barriers and a passage, the intensity of distracting events could be reduced. Third, the behavior room was large enough to enable multiple monkeys to enter and interact with the behavior station and each other with potential use in experiments of a social nature. Finally, we believe that our approach can be a practical blueprint for other monkey facilities who wish to implement an enriched living and behavior environment in smaller spaces. To this end, we provide a detailed description and specifications of various architectural, electrical, and mechanical components in our setup.

We commissioned a facility meeting our requirements which can house a small number of animals (3-6). This facility consisted of a naturalistic living room connected to a behavior room (with the behavior station) through a narrow but room-height passageway (Figure 1). Areas that were accessible by monkeys were separated from human accessed areas by either a solid laminate or toughened glass and stainless-steel mesh partition (Figure 1A).

The naturalistic living room was designed with the help of professional architects (Opus Architects & Vitana Projects) using guidelines developed for NHP facilities (Röder and Timmermans, 2002; Buchanan-Smith et al., 2004; Joint Working Group on Refinement, 2009). We incorporated ample opportunities for the monkeys to interact with the environment and used natural materials wherever possible. We provided two perches at above 2m elevation made of wooden beams on a stainless-steel frame (Figure 1B, 1C top), repurposed tree trunks as benches, and a dead tree as a naturalistic feature for climbing and perching. Cotton ropes were hung from the taller elements for swinging and playing. We also included a stainless steel pendulum swing for playing.

To prevent monkeys from tampering with and ensure their safety, all electrical components like roof lights and closed-circuit television (CCTV) cameras were enclosed with stainless-steel and toughened glass enclosures (Figure 1C, bottom). None of the structural and mechanical elements had sharp or pointed corners or edges. This room as well as other monkey accessed rooms described below were provided with a constantly replenished fresh air supply and exhaust ventilation. To keep unpleasant odors under control and to provide foraging opportunities for the monkeys, the floor of the living room was covered with a layer of absorbent bedding (dried husk and/or wood shavings) that was replaced every few days.

Compared to the older living area for monkeys (stainless steel mesh cages), the naturalistic living room designed by us is much more spacious (24 times the volume of a typical 1×1×2 m cage) and includes a large window for natural light. The living room is designed for easy removal and addition of features (all features are fixed with bolts and nuts), thus opening avenues for continuous improvement in enrichment conditions for the monkeys. The enriched living room was effective in engaging the animals as observed from heatmaps of their residence durations. Figure 1D shows activity in a 7 min period, both with and without the presence of humans (in the human interaction area). The animals heavily interacted with enrichment features leading to an observable improvement in their behavioral and social well-being.

From the living room, monkeys can approach the behavior room containing the behavior station (Figure 1I, touchscreen monitor for visual tasks and response collection) through a passageway between the living room and the behavior room (Figure 1H). The passageway is divided into two parts, the holding room and squeeze room (Figure 1E and 1F). The holding room is adjacent to the living room and is designed to be employed when isolating an animal as and when the need arises. A log bench was provided as enrichment in the holding room along with windows.

In between the holding and behavior rooms is the squeeze room. In the squeeze room, the back wall can be pulled towards the front to constrict the position and movement of the monkeys for routine tasks like intravenous injections, measurement of body temperatures, close inspection of a monkey by a qualified veterinarian, etc. The back wall is attached to grab bars in the human interaction room (to push and pull it) and a ratchet system (Figure 1G top shows the handle to engage and disengage ratchet) to prevent the monkey from pushing back. This enables a single person to squeeze and maintain the position of the back wall without applying continuous force, allowing the person to focus on interacting with the animal and maximize its comfort.

All monkey-accessible rooms were separated by sliding doors (Figure 1G bottom for latch mechanism) that can be locked to restrict a monkey to any given room. Ideally, all the sliding doors would be left open and monkeys move freely across these rooms. We also considered the need to bring the monkeys out of the facility for maintenance or relocation purposes and integrated trap doors (Figure 1E) in the doors of the living, holding and squeeze rooms. Onto these trap doors a transfer cage or even traditional monkey chairs may be positioned to move monkeys out of the facility for weighing, relocation to the colony, introduce new monkeys into the facility, etc. We successfully employed positive reinforcement training methods (Prescott et al., 2005; Rennie and Buchanan-Smith, 2006) to train our monkeys to come into the chair or transfer cage without having to employ the squeeze mechanism and trained them to move across the various rooms without hesitation.

Finally, the behavior room hosts the behavior station on the wall separating it from the control room (Figure 1A). The behavior station consists of a touchscreen monitor and juice delivery arm (Figure 1I) and is mounted on stainless steel channels which allow the panels and hence the behavior station to be modular. These panels can be repositioned or swapped with modified panels, so any change can be accommodated. The rest of the behavior room is also covered in these panels (except the floor but affixed to walls unlike the behavior station panels) which are layers of high-pressure laminate with a thin copper sheet sandwiched in the middle (Figure 1l and Figure S1A). This was done to insulate the behavior room from electromagnetic radiation from electrical and electronic circuitry present in the building and adjacent experimenter control room that housed all our computer systems. The resulting reduction in the magnitude of electromagnetic radiation across a wide frequency band (0-2KHz) inside the behavior room (Figure S1B) provides an ideal environment to collect neurophysiological recordings from the monkey’s brain using implantable electrodes and wireless recordings.

### Touchscreen workstation

Along with enriching their living conditions, we wanted to enrich the monkey’s experience when interacting with the behavior station. To this end, we decided to go ahead with wireless neurophysiological recording which meant that we do not need to restrict the monkey in front of the behavior station anymore. This decision opened us to opportunities to train monkeys in complex tasks that we can employ to understand the representation of objects in high-level visual areas of the monkeys’ brain. Conventionally, response choices to tasks are collected from chair restrained monkeys by either training them to make saccades with their eyes or by moving a joystick/bar provided near their hands. Instead, we chose to collect responses via a touchscreen monitor.

We affixed a commercial grade 15” capacitive touchscreen monitor from Elo Touch Solutions Inc. (1593L RevB) to the modular panels at the behavior station (Figure 2A, 2B). The height of the monitor from the floor was chosen such that the center of the screen lined up with the eye-height of a monkey sitting on the floor in front of the behavior station. This display supported a resolution of 1366 pixels by 768 pixels with a refresh rate of 60Hz and the polling rate of the integrated projected-capacitive touch panel was ~100Hz. The stimulus monitor and a second identical monitor (backup/observation unit located in the control room) were connected to a computer running the NIMH MonkeyLogic (NIMH ML, Hwang et al., 2019) experiment control software (running on MATLAB 2019a). Digital input and output of signals was facilitated by a National Instruments PCI-6503 card and BNC-2110 connector box combination (DIOxBNC).

Above and below the monitor on the behavior station were two acrylic window openings (17.7 cm tall by 22.8 cm wide) which enabled us to position a commercial infrared eye-tracker (ISCAN Inc., ETL 300HD, details below) below the monitor and a CCTV camera (frame sync-pulse recorded in NIMH ML through DIOxBNC) above the monitor (Figure 2A and 2B). We fine-tuned the relative placement of our binocular eye-tracker and synchronized network cameras to observe fine-grained eye movements as well as head and body pose of our animals as they perform different visual matching tasks (Figure 2C). A photodiode was placed on the touchscreen (Figure 2B) to measure the exact image onset times.

Because monkeys had to sip juice from the reward arm, this itself led to fairly stable head position during the task. To further stabilize the head, we designed modular head frames at the top of the reward arm (Atatri Inc), onto which monkeys voluntarily rested their heads while performing tasks (Figure S1D). We formed a variety of restraint shapes with stainless-steel based on 3D scans of our monkeys with progressively increasing levels of restriction (left to right, Figure S1D). Positioning their heads within the head restraint was not a challenge for the monkeys and they habituated to it within tens of trials. We also iterated on the structure of the reward arm, head restraint and fabricated custom attachments (hand grill, Figure 2A) that allow the monkey to comfortably grip at multiple locations with its feet and with the free hand and this in turn greatly reduced animal movement while providing naturalistic affordances on the reward arm (Figure 1H, right most panel).

The reward for performing the task correctly was provided to the monkey as juice drops delivered at the tip of a custom reward delivery arm (Figure 2A, 2B, and Supplementary Figure 1C). This reward arm was a 1” width hollow square section stainless steel tube. Concealed within it are two thin stainless-steel pipes – a juice pipe for delivering the juice to the monkey and a drainpipe to collect any remaining juice dripping from the juice pipe. The juice was delivered using a generic peristaltic pump on the pipe connecting the juice bottle to the end of the juice pipe in the control room. This pump was controlled by a custom voltage-dependent current driver circuit printed to a PCB (Figure S1E-F) which in turn is controlled through a digital signal from NIMH MonkeyLogic via the DIOxBNC board. The reward arm was mounted on a linear guide which allowed us to adjust the distance of juice pipe tip (near monkeys’ mouth) and the touchscreen. As a result, we can passively ensure the monkey sat at a distance that enables it to give touch response without having to stretch their arms and gave a good field of view of the monkeys’ face and body for the cameras.

### Eye-tracking in unrestrained animals

Eye movements were recorded using a customized small form factor ETL 300HD eye tracker from ISCAN Inc., USA with optical attachments that enabled eye tracking at close quarters. The eye-tracker primarily consisted of an infrared monochrome video capture system that we oriented to get a field of view that covered both eyes of the animal when its mouth was positioned at the juice spout and the animal was in position to do trials. The ISCAN system offers a parameterizable eye-gate, which is in effect a rectangular aperture in the monochrome camera’s field of view and restricts the search space of the pupil and eye-glint search routines in the ISCAN software algorithm. The pupil and eye-glint search are based on the area (minimum number of pixels) and intensity-based thresholds that can be manipulated using interactive sliders in ISCAN’s DQW software. We modeled the raw eye-gaze signal as the horizontal and vertical signed difference between centroids of the detected pupil and eye-glint regions of interest. The raw eye signal was communicated in real time to the computer running NIMH ML through the DIOxBNC analog cables. This raw eye-signal was read into the NIMH ML software and got rendered in real time onto another monitor that displayed a copy of the visual stimuli shown on the monkey touchscreen, while the monkeys performed touch-based visual tasks.

NIMH ML has a feature to display visual cues at selected locations on a uniform grid that the monkey can either touch or look at and obtain the liquid reward. We trained our monkeys to look at and then touch these visual cues. Since monkeys typically make an eye movement while initiating and performing the reach and touch, we exploited this to first center the raw eye signal with respect to the center of the screen and subsequently obtain a coarse scaling factor between changes in the raw eye signal and corresponding changes in the on-screen location. In this manner, we obtained a rough offset and scaling factor that maps the raw eye gaze signal with the on-screen locations of the monkey touch screen. We then ran calibration trials where 5 rectangular fixation cues were presented at pseudo-randomly chosen locations. The animal had to look at each fixation cue as and when it was shown, all the while maintaining hold on a button on the right extreme portion of the screen. The animal received a liquid reward at the end of a complete cycle of fixation cues for correctly maintaining fixation throughout the trials. These calibration trials provided us with pairs of raw eye-gaze (x, y) observations that corresponded to known locations on the touch screen. Although our monkeys were highly trained, the unconstrained nature of our experiment setup and session-wise and trial-wise noise across trials and sessions, requires us to use robust model fit procedures that learn the optimal linear mapping between raw eye-gaze x, y data and the on-screen locations, while identifying and rejecting outliers that are noisy of a result of saccades. We used the *fitlm* function provided in MATLAB with robust fit options turned on for this purpose.

We find that this method gives a satisfactory quality of raw eye-data to touch screen location mapping. We used these session-wise calibration models to transform eye-data if a higher degree of accuracy was required than what is provided by the initial coarse offset and scaling of the eye-signal that we manually perform in the beginning of each trial. We find that even the coarse centering and scaling of raw eye-data is sufficient for gaze-contingent paradigms where animals are required to look at and hold fixation at screen center, while stimuli are flashed in passive fixation tasks or while the animals maintain fixation at screen center while sample and test images are shown in succession in the same/different tasks.

### Social training of naïve monkeys

The aim of the social training was to investigate whether a naive animal can learn the same-different matching task faster (than TAT) by observing trained monkeys performing the task. Here the naive animal had never interacted with touch screen before and learnt the task from scratch. Learning happened by trial and error, observation and cooperation.

#### Animals

We introduced M2 to the behavior station with M1 and M2 for social training. M2 was group-housed with M1 and M3 from 9 months before start of social sessions, so their social hierarchy was observed to be M1 > M2 > M3.

#### Stimuli

A set of 100 images of unique natural objects were used as stimuli. On Day 21 and Day 29, a new set of 50 images of unique natural objects were used to test the performance. All stimuli were presented after conversion to grayscale and the longer dimension of the images was always equated to 5.5° visual angle. Images were taken from the BOSS v 2.0 stimuli set (Brodeur et al., 2010, 2014).

#### Training

Temporal same-different task (stage 10 of TAT, Figure S2) was chosen for the social training sessions. Unlike TAT where an animal progressively attempts stages of the task until it is proficient in the full task, in social training sessions we investigated how a naïve monkey might learn the full task in the presence of trained peers (M1 and M3). Crucially, M2 can only get access to juice reward by responding when choice buttons are presented at the latter half of the trial.

Sessions were held on all mornings of the week except for Sundays and only if animals voluntarily moved to the behavior room (animals were herded two at a time through to behavior room, closing partition doors behind them). If any animal did not come for a particular session, it was supplemented with 50 ml of water. For instance, M3 did not come on Day 3 and Day 7; for these sessions, M2 was introduced alone into the behaviour room. Weight was monitored continuously as described earlier.

On each social session, we introduced M2 along with M1 (its superior in social rank) for 15-20 minutes or until M1 performed ~400 correct trials or 80 ml of juice. On the same day, we also introduced M2 with M3 (its subordinate in rank) for 45 minutes or until M2 received 60 ml of juice. Interestingly for few trials M2 and M3 cooperated (day 4: 35 trials, day 5: 14 trials, day 8: 96 trials and day 9: 10 trials; Video S5 and Figure S4). M2-M3 session was for 45 minutes or until M2 received 50-60ml of juice, whichever was earlier. Video recordings of both the sessions were done for subsequent coding of distinct behavioural episodes in these sessions.

Previous studies have established that animal learns more from peer’s mistake (than from peer’s success) and from own success (than own mistake) (Monfardini et al., 2012, 2017; Isbaine et al., 2015; Ferrucci et al., 2019). In a two-choice task, error reduces the preference of the choice made by the animal (Monfardini et al., 2017). In our case, the error signal is generated from multiple sources: breaking hold maintenance, incorrect response and no response. We felt that maintenance of hold, before the sample is shown is not crucial to task performance. Hence, we choose to make the task much easier and reduce errors by reducing the initial hold time down to 100ms which reduced the hold maintenance time to 700ms from 1.1 second. When the monkey started to get reward on 50% of responded trials, we increase the initial hold time to be 300ms on day 16 and 500ms on day 17. After that the hold was 500ms throughout the training. We modified inter-trial intervals (for correct and incorrect responses) and reward amount to keep M2 motivated to learn the task.

On Day 5, for few trials M2 was able to maintain the hold till the response buttons appeared. Then he dragged his hand below and touched the “different” response button (which was positioned at the bottom of hold button). He was able to obtain a reward on 50% of the responded trials using this biased strategy. To discourage him from choosing only “different” button, on Day 6, we enabled immediate repeat of incorrect trials, so that an error trial was repeated immediately until he made a correct response. From Days 7-9, immediate repeat of error trial was disabled but on Day 10 we re-enabled immediate repeat of error trials to remove response bias. Once M2’s overall accuracy on responded trials (including immediate repeat of error trials) reached 80% (Day 20) we disabled immediate repeat.

#### Social session analyses

Since two monkeys were in the behaviour room during social sessions, we first identified which trial was done by which monkey by manually annotating the CCTV videos. Then for each monkey, we calculated accuracy on responded trials as a percentage of correct trials out of responded trials (Figure 4B). Accuracy could be of two types: First chance accuracy was calculated on all responded trials without including immediate repeat of error trials. Second-chance accuracy was calculated only on immediate repeat of error trials (after making an error, there were a stretch of same trial repeating, until the monkey made a correct response). For M1, repeat of error trials were not activated, and in case of M3, days when he did the task (day 4, 5, 8 and 9) immediate repeat of error trials were disabled. For M2-M3 session, we calculated percentage of trial initiated by M2 and percentage of trial initiated by M3, on total trials of that session (Figure 4B inset).

To understand the learning stages of M2 (Figure 4C), we calculated touching accuracy (percentage of total trial where M2 initiated the trial by touching), response accuracy (percentage of total trials in which M2 made a response) and correct response accuracy (percentage of total trials where M2 made a correct response). These three accuracies were calculated on total trials attempted by M2 alone (excluding the trials performed by M3).

## AUTHOR CONTRIBUTIONS

All authors were involved in the entire study throughout, but took individual responsibility for specific aspects as follows. GJ, HK, ZK and SPA conceptualised the new lab. GJ, HK, ZK and SPA worked with Opus Architects to finalize the design with Opus Architects, and GJ & SPA worked with Vitana Projects for the implementation. HK, JD, TC and SPA conceptualized the reward arm and head restraints and oversaw fabrication. HK, JD and TC oversaw and coordinated steel-works, and prototyping and testing of fabricated products. HK, TC and SPA worked with ISCAN in customizing the head-free eye-tracker. ZK and SPA designed, identified and procured equipment required for behavioural and neural data monitoring and recording. TC and ZK performed system integration and testing. GJ, HK, JD, TC and ZK wrote MATLAB-based codes for behavioral training. JD and ZK performed shield testing. GJ and TC oversaw design and fabrication of juice delivery systems. GJ, HK, JD and TC worked on all aspects of monkey training. GJ, HK, JD, TC and SPA wrote the manuscript with inputs from ZK.

## ACKNOWLEDGMENTS

We thank Sujay Ghorpadkar (Opus Architects), Anagha Ghorpadkar (Vitana Projects), Rikki Razdan & Alan Kielar (ISCAN), Assad & Mahadeva Rao (Fabricators), Ragav (Atatri) for their excellent services in developing all custom components. We thank Mr. V Ramesh (Officer in-charge) and Ravi & Ashok (workers) from the Primate Research Laboratory (PRL) for their excellent animal husbandry and facility maintenance.

## Data Availability

All the data and codes required to reproduce the results are publicly available at https://osf.io/5764q/

## SUPPLEMENTARY MATERIAL

### SECTION S1. CUSTOM DESIGN ELEMENTS

**Figure S1:**
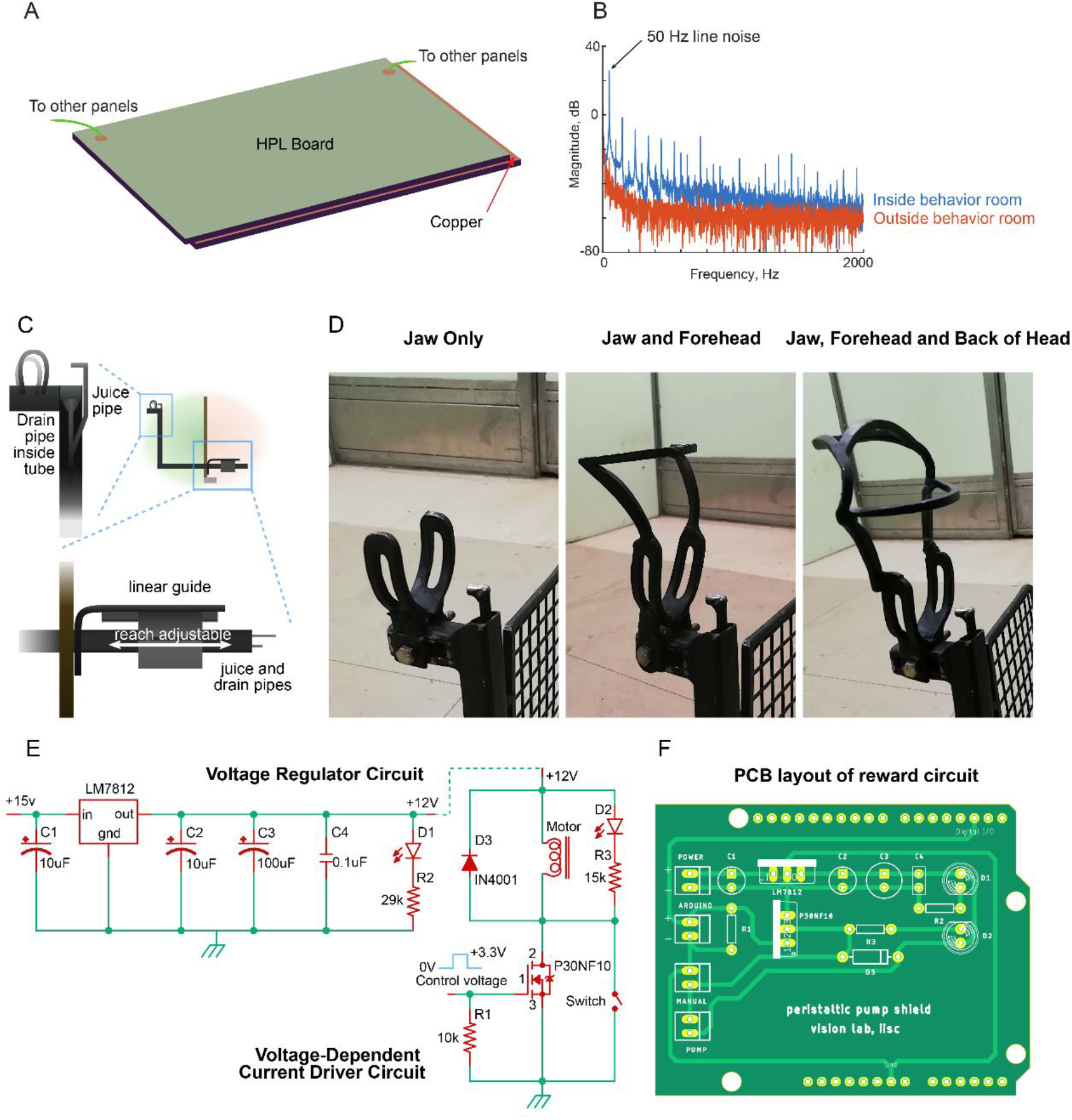
Custom design elements in the behavior room. (A) Schematic of the copper sheet sandwiched between layers of high-pressure laminate panels. These panels are installed on the walls and roof of the behavior room and electrically connected so as to form a closed circuit to block external radio frequency noise. (B) Power spectrum (in dB) of noise recorded from the behavior room with shielding (*red*) and the control room without shielding (*blue*). The copper sandwiched panels in the behavior room and all stainless steel supporting frames were connected electrically to the ground of the pre-amplifier (Plexon Inc). Signals were recorded at 40 kHz for 1 s using a 24-ch Uprobe electrode floating in air connected to a 32-channel data acquisition system (Plexon Inc). (C) Schematic of juice reward arm. At top right, a close-up view of the spout portion of the juice reward arm showing how the juice pipe and drain-pipe are concealed within a tubular stainless-steel pipe. This prevents monkeys licking any run-off juice or from tampering with the thin steel juice pipe itself. Bottom close-up shows how the juice reward arm can be moved into and out of the behavior room to accommodate the monkey’s hand reach (using a lockable linear guide). (D) Photographs of three head frames with increasing levels of restraint (left to right). Each restraint is made from stainless steel rods bent to match the typical shape of the monkey head (obtained using 3D scanning). (E) Circuit diagrams of the voltage regulator (*left*) and voltage-dependent current driver circuits (*right*) that are part of the reward system. (F) Layout of the printed circuit board (with the voltage regulator and voltage-dependent current driver circuits from panel E). This circuit board powers a peristaltic dosing pump to push juice into the juice pipe.

### SECTION S2. TAILORED AUTOMATED TRAINING (TAT)

Here we describe the Tailored Automated Training (TAT) paradigm we used to train naïve monkeys to perform a same-different task.

#### RESULTS

The complete trajectory of training for both M1 & M3 is depicted in Figure S2 and are summarized below.

Stage 1 was the touch stage: here monkeys had to touch a green square that appeared on the screen upon which it received a juice reward. Both monkeys cleared this stage easily (Figure S2).

In Stage 2, monkeys had to hold their fingers on the green hold cue for increasing durations (100 to 3000 ms). The hold time was small initially (100 ms) so that monkeys would be rewarded for accidentally long touches and start to hold for longer periods. We trained monkeys to hold for longer periods (3 s) since this would be the hold time required eventually for the same-different task. Towards the end of this stage, we began to flash successive stimuli (upto 8 stimuli with 200 ms on and off) at the center of the screen while the monkey continued to maintain hold. Both monkeys took about two weeks to clear this stage (15 sessions for M1 to reach 2.6 s, 13 sessions for M2 to reach 3 s; Figure S2).

From Stage 3 onwards, monkeys started seeing a simplified version of the same-different task. Here we tried many failed variations before eventually succeeding. In Stage 3, they maintained hold for 500 ms, after which a sample image was shown for 400 ms, followed by a blank screen for 400 ms. After this a test image was shown at the center and the hold cue was removed, and a single choice button appeared either above (for SAME trial) or below (for DIFFERENT trial). To simplify learning, we used only 2 images resulting in 4 possible trials (either image as sample x either image as test). Monkeys had to release hold and touch the choice button within a specified period of time. Once monkeys learned this basic structure, we reasoned that reducing this choice time would force them to learn other cues to predict the choice button (i.e. the sample being same/different from test). However this strategy did not work and we discarded this strategy after 16 sessions (Figure S2).

In Stage 4, we introduced both choice buttons but the wrong choice button had a lower intensity to facilitate the choice. Both monkeys quickly learned to select the brighter choice button. Here our strategy was to reduce the brightness difference to zero, thereby forcing the animals to learn the same-different contingency. Here too, monkeys kept learning to discriminate finer and finer brightness differences but failed to generalize to the zero brightness conditions. We discarded this strategy after 13 sessions (Figure S2).

In Stage 5, we tried several alternate strategies. These included immediate repeat of error trials (thereby allowing the monkeys to switch to the correct choice button), overlay of the image pair on the correct choice button (to facilitate the association of the image pair at the center with the choice buttons). While monkeys learned these associations correctly, they still did not generalize when these conditions were removed. On closer inspection, we observed that this was because they were looking only at the response button and not at the sample and test images. We discarded this strategy after 13 sessions (Figure S2).

In Stage 6, we further simplified the task by keeping the sample image identical in all trials, and varying only the test image (i.e. AA vs AB trials). We also simplified the task by showing the sample throughout, and then displaying the test image alongside the sample after a brief delay to facilitate comparison. We initially overlaid the image pair on the correct response button and eventually removed it based on performance. Monkeys cleared this level easily, and encouraged by this success, we introduced pairs of trials with new image pairs. In each level the old/learned pairs had no overlay (these were 50% of the trials) and the new pairs had overlay (these were the remaining 50%). In this manner, we introduced 20 image pairs made from 20 unique images. Note that clearing this stage means that monkeys might have learned the full same-different concept or alternatively learned to associate specific test images to the “SAME” or “DIFFERENT” choice buttons. Monkeys cleared this stage in 8 sessions (Figure S2).

In Stage 7, we attempted to nudge the monkeys towards a full same-different task. Here we used 8 new images such that the test image was always the same in a given pair but the sample image varied (i.e. AA vs BA trials). Monkeys cleared this stage in 3 sessions (Figure S2).

In Stage 8, we combined the trials from Stages 6 & 7 in equal proportion (8 image pairs each). Monkeys cleared this stage in 1 session (Figure S2). However, it is still possible that they were doing this task by remembering sample or test associations with the corresponding choice buttons.

In Stage 9, we introduced all possible image pairs possible from 20 new images along with the previously learned image pairs and gradually reduced the proportion of the learned pairs. Both monkeys cleared stage easily (6 sessions for M1, 5 sessions for M2), suggesting that they learned the concept of same-different. We further confirmed this by testing them on 100 new images, where sample and test images were chosen randomly from the ^100^C_2_ = 4,950 possible sample-test pairs. Monkeys cleared this stage in 13 sessions (Figure S2).

In Stage 10, we transitioned to a temporal same-different task by reducing the temporal overlap between sample and test images, introducing a brief delay period and then gradually moving the test image to the same position as the sample. Monkeys easily cleared this stage in 4 sessions (Figure S2).

**Figure S2:**
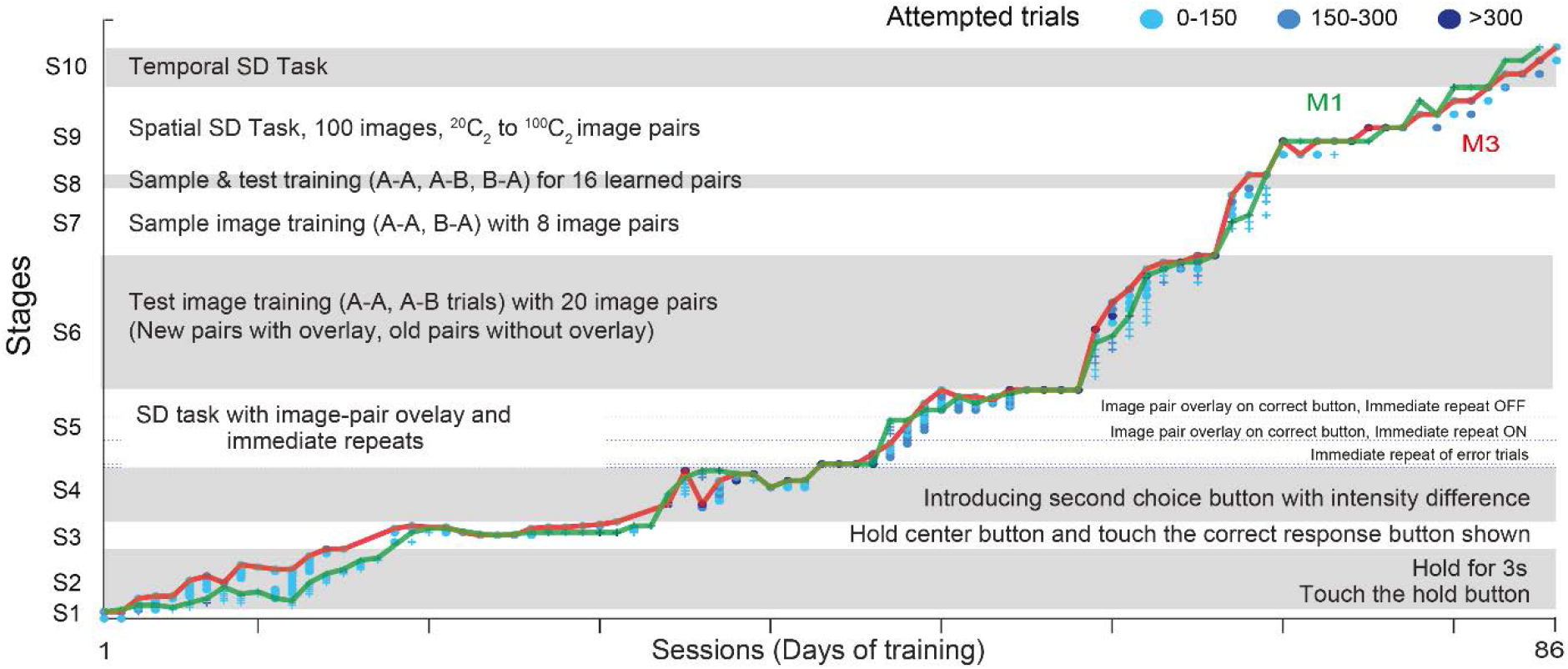
Tailored Automated training (TAT) on Same-Different task. The plot shows the progression of animals M1 and M3 through the ten stages of TAT. Each stage is further divided into levels with symbols corresponding to each monkey (plus for M1, circles for M3) and color indicating the number of trials attempted (0-150 trials: *light blue*, 150-300 trials: *cyan*, >300 trials: *dark blue*). The lines indicate the maximum level reached by each animals in a given sessions (M1: *green*, M3: *red*).

#### METHODS

##### Animals

M1 and M3 participated in the Tailored Automated Training. The animals were each provided a 45-minute period of access (session) to the behavior station with no fixed order of access. Training was conducted only if animals voluntarily moved to the behavior room. Animals were moved one at a time through to behavior room, closing partition doors behind them. If the animal was not willing to go forward to the behavior room, training was not done on that day and the animal was supplemented with 50 ml of water later in the day. Weight of the animals were checked twice a week and if any sudden drop in weight was measured the animal was given time to recover (by removing water restriction and pausing training).

##### Stimuli

For TAT, stimuli were selected from the Hemera Objects Database and consisted of natural and man-made objects with a black background to match the screen background.

##### Training

The aim of the TAT was to teach monkeys the temporal same-different matching tasks (SD task), a schematic of which is shown in figure 3A. We employed TAT as a proof of concept to show that it is possible to achieve unsupervised training for animals on a complex same-different (SD) matching task. We automated the training by dividing the SD task into sub-tasks (stages) with further levels within each stage to titrate task difficulty. Animals progressed to successive levels and stages based on their performance (when accuracy on the last 50 attempted trials within a session was greater than 80%). Like recent automated training paradigms (Berger et al., 2018), we provided an opportunity to go down a level, if the animal performed poorly but we ultimately moved to a more stringent level progression where the animals were not allowed to slide back to an earlier level/stage. We started from a lower level only when the training was resumed after a long break, due to unavoidable circumstances like equipment failure or issues related to animal health. Overall, we find that the rate of learning depends on animal’s underlying learning capability and the design of the automated training regime. Hence to achieve fastest learning rates, we optimized the level-wise difficulty of the automated design.

In general, the progression of task difficulty across levels and stages was selected such the animal could always perform the task at above-chance performance. Although we set out to train animals using a completely automated pipeline, we also wanted to ensure that both our naive animals could complete the learning process in full without drop out as is common in many automated regimes (Calapai et al., 2017; Tulip et al., 2017; Berger et al., 2018). We implemented a pragmatic approach, to intervene and tailor the training parameters at particularly difficult stages for so as to avoid the monkey dropping out of the training process entirely.

The SD task was divided into ten conceptual stages. A single parameter was varied across levels within a stage. The smallest unit of the TAT is a trial, but composition of each trial is dependent on the current level. Each trial started with the presentation of trial initiation button and trials were separated by a variable inter-trial interval (ITI). The duration of ITI depends on the outcome of the current trial (500 ms for correct trials; 2000 ms for incorrect trials). Provision was made to change some parameters quickly without aborting the experiment. The ITI and reward per trial were adjusted within a session based on animal’s performance. We increased ITI to give another level of feedback when animals were showing very high response bias by pressing only one button or when the animals were satisfied with 50 percent chance performance.

Liquid juice reward was delivered after every correct trial. We started each session with 0.2 ml of juice reward per trial. Juice reward was increased for consistent behavior but never decreased within a session. The motive behind increasing the reward was to keep the motivation high when learning a new task as any kind of error done by the animal aborts the trial. Monkeys got two distinct audio feedback tones: a high-pitched tone for correct response and a low-pitched tone for incorrect responses (including uninitiated, aborted or no response trials).

#### TAT Stages

##### Stage-1 (Touch)

A green button (square) was presented on the touch screen where monkey had to touch for reward. Any touch outside was considered as error. There were two levels in this stage (Button size: 200 x 600 pixels in level 1.1 and 200 x 200 pixels in level 1.2). Center of the buttons were same as the that of the hold button in Figure 3A.

##### Stage 2 (Hold)

The hold button was presented and monkeys had to touch and maintain the touch within the button area until it was removed. Any touch outside the hold button was considered an error. There were thirty levels in this stage, in which hold time varied from 100 ms to 3 s in equally spaced intervals. M3 cleared all the levels but M1 was trained only up to a hold time of 2.6 s.

##### Stage 3 (1-Response Button)

A temporal same different task with only correct choice button was presented. Choice buttons were green colored squares and were presented above and below the hold button for same and different choices respectively. Image presentation sequence was same as that shown in Figure 3A. We had a wait to hold time for initiating the trial as 8000 ms, pre-sample delay time of 500 ms, sample-on time of 400 ms and post-sample delay of 400 ms. We reduced the time to respond in this level from 5 s to 400 ms in several steps (in 1000 ms steps till 1s, 100 ms steps till 500 ms and 50 ms steps till 400 ms). Four image pairs formed from two images were used to construct the same different task.

##### Stage 4 (2-Response Buttons)

In this stage the wrong choice button (also of similar dimensions and color to the hold button) was also displayed with brightness that increased from 0 to the maximum intensity (same as the correct choice button). This is a full temporal same different task with an intensity difference between correct and wrong choice buttons. Wrong button was introduced in ten steps with brightness scaled relative to the maximum intensity (scaling factor for each level: 0.2, 0.4, 0.5, 0.8, 0.85, 0.90, 0.925, 0.95, 0.975, 1). A scaling factor of 1 meant that there was no intensity difference between the choice buttons, and the monkey would have to use the visual cues (sample & test images) to perform the task. Time to respond was 800ms and all other task parameters are same as stage 3.

##### Stage-5 (Ad-hoc Strategies)

We introduced two new strategies (Immediate Repeat and Overlay) to facilitate same-different training. With the immediate repeat strategy, for every wrong trial, we repeated the same trial again with a lower reward (0.1 ml) for correct response. This allowed the animal to switch its response upon making an error. In the overlay strategy, we presented images of sample and test side by side blended on the correct choice button (blended image = α*image + (1-α)*choice button), where α is a fraction between 0 and 1. We started the first level of this stage by giving three kinds of additional information (Button intensity difference, Immediate Repeat and Overlay) to identify the correct response. As the levels progressed, we removed the cues slowly. First, we removed button intensity difference in 6 levels (scaling factor of wrong button intensity in each level: 0.2, 0.3, 0.5, 0.7, 0.9, 1). Second, we removed the overlay cue in 15 levels. (Blending factor α: 0.5, 0.4, 0.3, 0.2, 0.15, 0.1, 0.09, 0.08, 0.07, 0.06, 0.05, 0.04, 0.03, 0.02, 0.01,0). We removed the immediate repeat of error when blend cue reached α = 0.06.

##### Stage-6 (Test Stimulus Association)

Stages 6, 7, 8 and 9 were based on a spatial version of the same-different task. In Stages 6 and 7, a new condition was introduced with overlay on correct response and this happened on 50% of trials in trial bag. The remaining trials were already learned conditions which were shown with no overlay. A level with overlay on correct response was repeated with a level without overlay. This spatial task differed from the temporal tasks in the position of the test image (shifted right or between sample and hold button) and sample ON time (sample image is presented till the trial ends). Each level introduced two new images through two specific image pairs (Images A and B are introduced through trials AA and AB). The trials only differed in the test image, so the monkey can do the task only by associating a test stimulus to the correct choice button. In all, we introduced 20 new images and 20 image pairs across levels. Since we were presenting newly introduced image pairs more often (ratio of new image pairs to learned image pairs is 1:1), the monkeys could reach 80% accuracy without attempting all learned image pairs. Hence, to check the monkey’s performance on all learned image pairs, we created the last level with all 20 image pairs presented equally likely without cue.

##### Stage-7 (Sample Stimulus Association)

In this stage we introduced image pairs formed from two images which differed in sample image (Images A and B are introduced through image pairs AA and BA but not AA and AB). In total we introduced 8 new image pairs formed from 8 images. All other experimental conditions were same as Stage-6

##### Stage-8 (Sample and Test Association)

Here we presented 16 image pairs selected from Stage-6 and Stage-7 together.

##### Stage-9 (Spatial same-different task)

All possible image pairs from 20 new images were introduced in this level and this was done along with learned pairs (ratio of new pairs is to learned pairs is 1:1 with new pairs shown with choice button overlay). In next level overlay was removed and in subsequent levels the proportion of new image pairs were increased (this was done in two levels: 75:25 and 100:0). We tested the generalization introducing two new set of images (number of images in these sets: 20 and 100) in next two levels.

##### Stage-10 (Temporal same-different task)

The task was switched to temporal from spatial SD task. In the first level we retained the sample image and test image location but we turned off the sample image before presenting the test image. There was no delay between sample and test. Next level, the sample and test were spatially overlapping and the delay between sample and test were zero. In the subsequent levels the delay between sample and test were increased in steps (50 ms, 100ms, 200ms).

### SECTION S2. ADDITIONAL ANALYSES OF SOCIAL LEARNING

**Figure S3:**
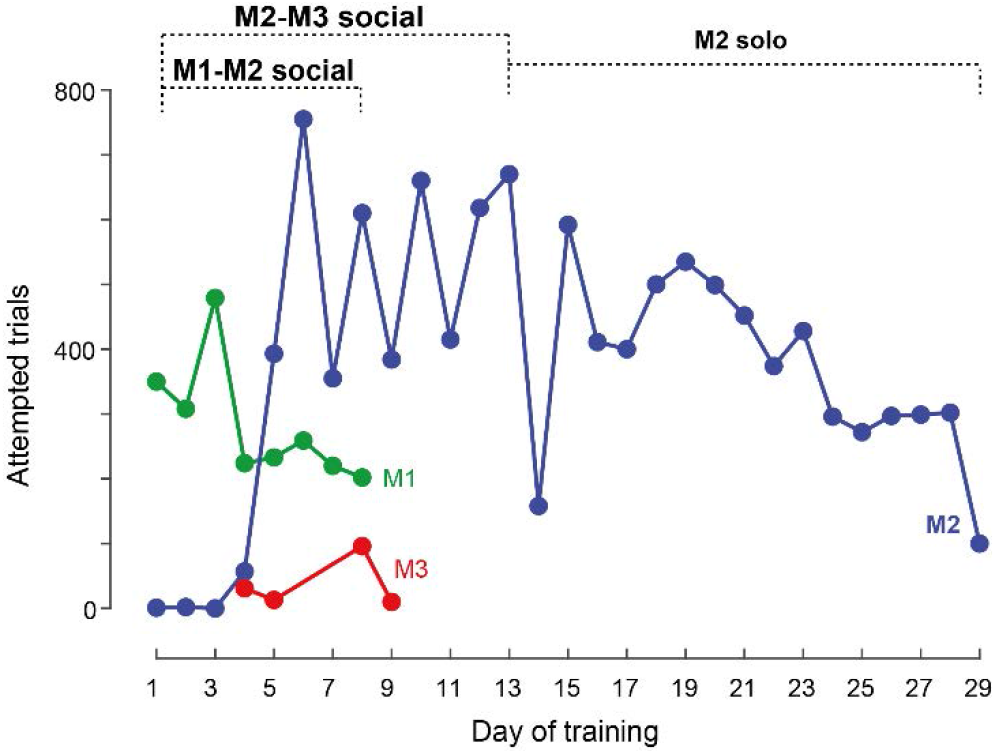
Daily trial summary of Social learning. Attempted trials during social training sessions (green-M1, blue-M2 and red-M3) across days.

**Figure S4:**
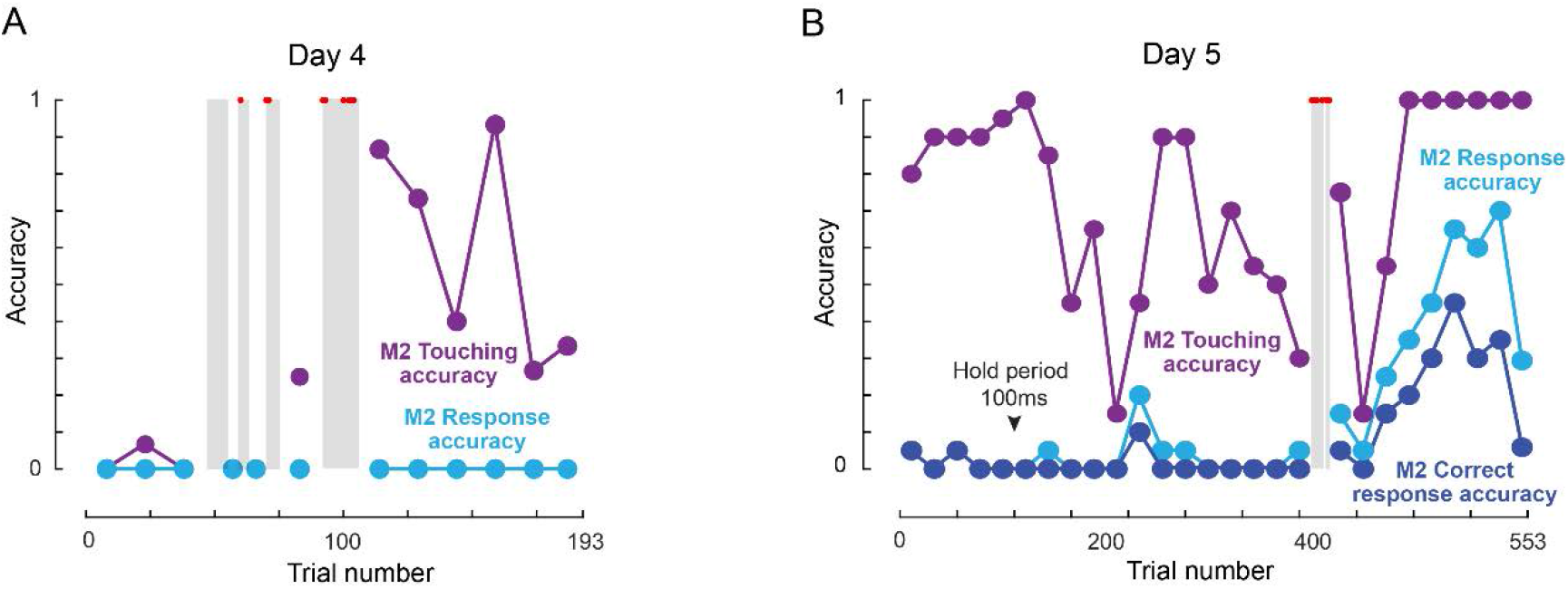
M2-M3 co-operation during social learning. (A) Day 4 M2-M3 session: Shaded regions are showing trials where M2 and M3 co-operated in the task (M3 performed the task and M2 got juice). Red dots in shaded region are showing correct trials. The whole session is divided into non-overlapping bins (bin size is 15 trials except in the shaded regions). Each dot represents accuracy calculated on the total trials in that bin. *Touching accuracy*: percentage of trials initiated by M2. *Response accuracy*: percentage of responded trial (correct or incorrect) out of total trials. (B) Day 5, M2-M3 session: *Correct response accuracy*: percentage of total trials in which M2 made a correct response. Here bin size is 20 trials. All other conventions are same as (A).

